# Safety Assessment of Intravenous Injection of Rat Embryonic Proteome Extract for *in-vivo* Regenerative Therapies

**DOI:** 10.1101/2023.10.30.563384

**Authors:** Siva Rama Prasad Darsi, Siva Kumar Kandula, Kala Kumar Bharani, Anil Kumar Pasupalati, Satyanarayana Swamy Cheekatla, Sujesh Kumar Darsi, Adi Reddy Kamireddy, Ram Reddy Barra, Ashok Kumar Devarasetti, Sreedhar Surampudi, Jaya Ram Raddy Singireddy

## Abstract

**Back ground:** Tissue differentiation and organogenesis are exclusively embryonic-phase activities, and these two activities are strictly under the control of the embryonic proteome, the developing embryo is a prime source of molecules for regenerative therapy. Many *ex-vivo* rudimentary organoid culture studies confirmed the inherent biological function of embryonic factors. Before using embryonic proteins for in vivo therapies, their safety should be confirmed first. Hence, we aimed for an *in vivo* study to inject rat embryonic proteome extract (EPE) through the intravenous route and investigate the impact on immunological, biochemical, and hematological parameters in the adult rats.

**Methods:** In this study, we isolated rat embryonic proteins from the 14^th^, 16th, and 19^th^ embryonic days by homogenization of embryos and isolated protein extracts through ultra-centrifugation. Six pairs of rats have been taken; six are allotted for control, and six are for the embryonic extract injection. The isolated embryonic protein extract was injected intravenously into the treatment group of rats and the normal saline into the control group. After the injections, blood samples are collected from both the treated and control groups to analyse immunological markers Il6, CRP, biochemical parameters creatinine, urea, sugar, proteins, albumin, globulin, and hematological parameters total leucocyte count, neutrophils, and lymphocyte percentage

**Result:** After the EPE injection, biochemical parameters, immunological markers, and hematological parameters were analyzed in both control and treated groups. All the above mentioned parameters are within normal limits. Statistical analyses were done using the t-test for unequal variance (p< 0.05). We observed no significant difference between the control and treated groups, so *in vivo* injections of embryonic protein extract is safe for the use of *in vivo* regenerative therapies.

**Conclusion:** Despite extensive *in vitro* studies confirmed the biological function of embryonic growth factors for organ differentiations and rudimentary organoids, but *in vivo* clinical applications are not yet started because of immunogenicity, heterogeneity, and tumorigeniety. In our study we injected EPE intravenously and showed that EPE is non-immunogenic, non-heterogeneic, and non-tumorigenic. This study concludes that EPE is safe for in vivo injection, so that further studies can be continued for intra-organ injections for organ regenerative therapy.

## Introduction

Regenerative medical research has stumbled into a new direction in the use of embryonic protein extracts (EPE). The dynamics of tissue differentiation, organogenesis, and growth are strictly operated through specific embryonic factors [1]. During the development of an embryo, tissue differentiation and organogenesis are manifested with embryonic differential expressions of the transcriptome [2] and proteome [3]. Proteomic analysis of a study has shown that 977 embryonic proteins are unique among 288 adult proteins. This determines that embryonic cell transcriptomes [4] differ from adult cell transcriptomes [5]. Thus, the embryo is a reservoir of different proteins that might be involved and can be used for regeneration and growth of damaged organs in the adult stage. Recent comparative in vitro studies have confirmed the embryonic proteins have regenerative function [6].

One of the prominent studies is the transformation of adult somatic cells into induced pluripotent stem cells by using the Yamanaka factors Oct3/4, Sox2, Klf4, and c-Myc [7]. All these factors are exclusively expressed on the 14th day of the rat embryo. One more in vitro study has confirmed that early nephrogenesis is due to epigenetic factors like Cbx1, Cbx3, Cbx5, and Trim28 isolated from developing kidneys [8]. Cell cultures, while supplemented with these factors, will be differentiated into ureteric buds and elongated into renal vesicles, forming comma– or S-shaped bodies. In another study, proteins from vesicles of amniotic fluid stem cells are injected through the intra-ventricular route; these injected cells are homed in the damaged kidney and exert a reno-protective function [9]. All these studies have paved the way for embryonic molecules’ application in the regeneration of organs.

Application of embryonic molecules is a recent advancement over stem cell therapy, as stem therapy has crucial limitations tumorgenicity[10], heterogeneity[11] and immunogenicity [12]. Tumorgenicity and heterogeneity are intact cellular functions, so in this study we have deployed homogenization of embryonic stem cell to decellurized [13], so that tumorgenicity and heterogeneity can be aimed resolve. Present study also confirmed that embryonic protein extract (EPE) does not activate immune cells, so it is safe for In vivo therapies. Total experimental work flow is elucidated in (Fig. 1). There are four major advantages to the application of embryonic proteins: non-tumorgenicity, non-heterogeneity, non-immunogenicity, and regenerative potency [14]. Very few studies and scant literature were observed regarding the determination of embryonic stem cell proteins compatibility during the development and repair of tissues and organs. The safety and compatibility of any therapeutic proteins including embryonic Protein extract injection was confirmed by biochemical, haematological, and immunological manifestations [15].

**Fig. 1.**
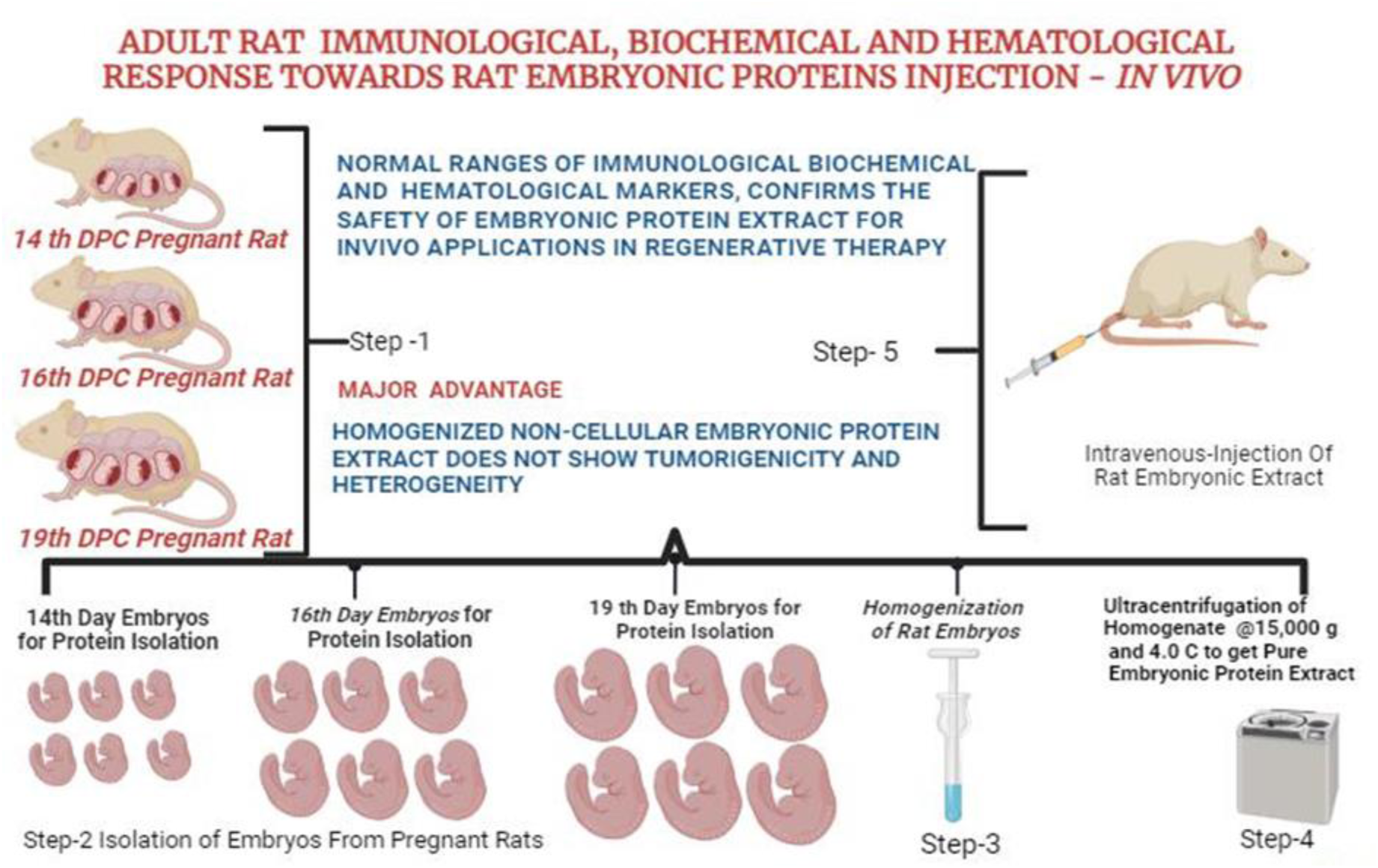
Total experimental Procedure presented in figure 1 in 5 different steps. **Step 1** includes selection of animals and caged for mating, conformation of mating through vaginal swab microscopy and vaginal plug formation. **Step 2** Scarification of 3 pregnant rats, on three different embryonic days 14^th^, 16^th^ and 19^th^ to isolate embryos. **Step 3** Homogenization of embryos of 14th, 16^th^ and 19^th^ embryonic days separately in EDTA 0.85 mg/ml. **Step 4** Protein extraction, after homogenization of 14^th^, 16^th^ and 19^th^ homogenates were, centrifuged out at 13,000× g and 4 °C for 30min. The supernatant was recentrifuged at 13,000× g and 4 °C for an additional 30min to get maximal purity. The pellet was discarded, and the final samples were was stored at −80 °C until use. **Step 5** Embryonic extract of different was given through intravenous injection (Image created using Biorender creator)

## Materials and Methods

### Selection of Rats

One pair of Sprague Dawley rats were provided with unrestricted access to food and water while being kept on a 12:12 h light cycle. All the animal were housed in isothermic environment as per animal ethical committee guidelines. To ensure biological replication, six pregnant rats were examined based on the cytology of vaginal smear microscopy and visible vaginal plug formation [16].

This first set, consisting of one female and one male, was allowed for mating to produce the first filial generation of six female and five male rats. Again, these five female and five male rats were arranged for mating to generate their second filial generation. In second filial generation 5 females are arranged for mating. Among these five, 3 pregnant rats are scarified for isolation of embryonic proteins in 14^th^ 16^th^ and 19^th^ day embryonic days respectively [17] and two rats continued full gestational period, so that 20 offspring’s of F2 generation nurtured. Out these 20 six males and six female rats selected for this study, 6 are for EPE injection and 6 are for control group, allocation of rats for this study is Illustrated (Fig-2).

**Fig. 2.**
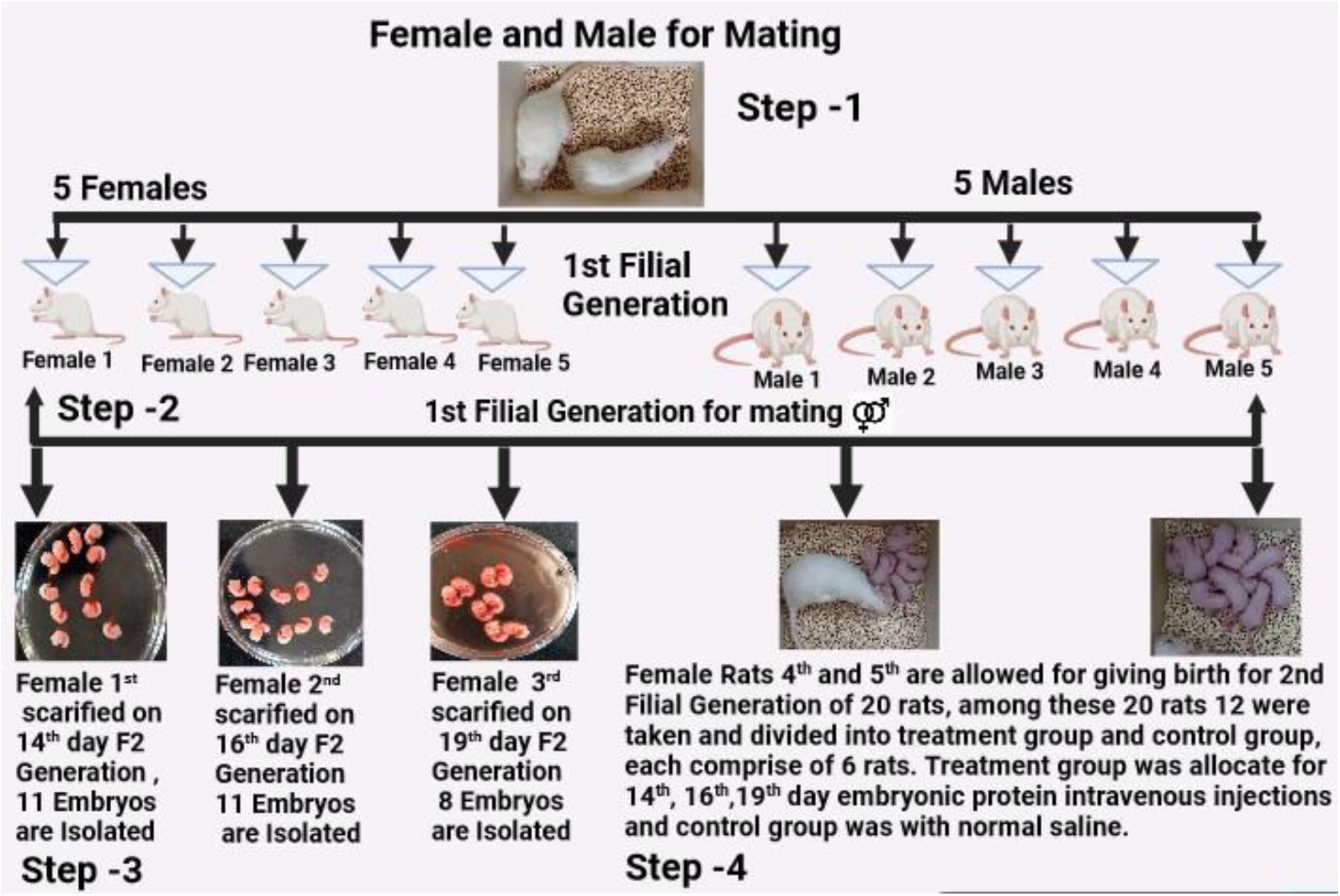
Selection and allocation of rats in different experimental steps**. Step 1** allocation for initial mating for or first filial generation**. Step 2** in-breading of f1 generation for f2 generation**. Step 3** Isolation of embryos on 14^th^, 16^th^ an d 19^th^ embryonic days of three pregnant**. Step 4** Twenty produced are in f2 generation, of these 20 rats 12 are divided in two groups, treatment group and control group for further study.

### Isolation and quantification of embryonic proteins

A total of 30 embryos were isolated on 14^th^, 16^th^ and 19^th^ embryonic days from 3 pregnant rats. A single embryo from each day was taken, washed six times with 10x PBS buffer and homogenized using an ultrasonicator in EDTA 0.85 mg/ml to inhibit the proteolytic enzyme activity [18]. The EPE used for *in-vivo* injections, and the entire process of extraction is performed under cold chain at 4^0^c. Remaining un-homogenized embryos are stored at –80^0^ c for future use. The isolated EPE were quantified on nanodrop spectrophotometer [19]. Protein concentrations of 14^th^, 16^th^ and 19^th^ embryonic days are 5.21 mg/mL, 12.19 mg/mL, and 24.1 mg/mL respectively.

### Polyacrylamide Gel Electrophoresis of EPE

The 10% polyacrylamide denaturing gel was used for the EPE protein separation. In the gel in the first well, 5 μL of the biorad reference standard was used, along with 5 EPE samples from days 14, 16, and 19 were loaded to the following wells and duplicate gel run was executed. After separation, gels were stained with Coomassie brilliant blue staining. The gel was destained and scanned, for further analysis, as shown in the (Fig. 3).

**Fig: 3.**
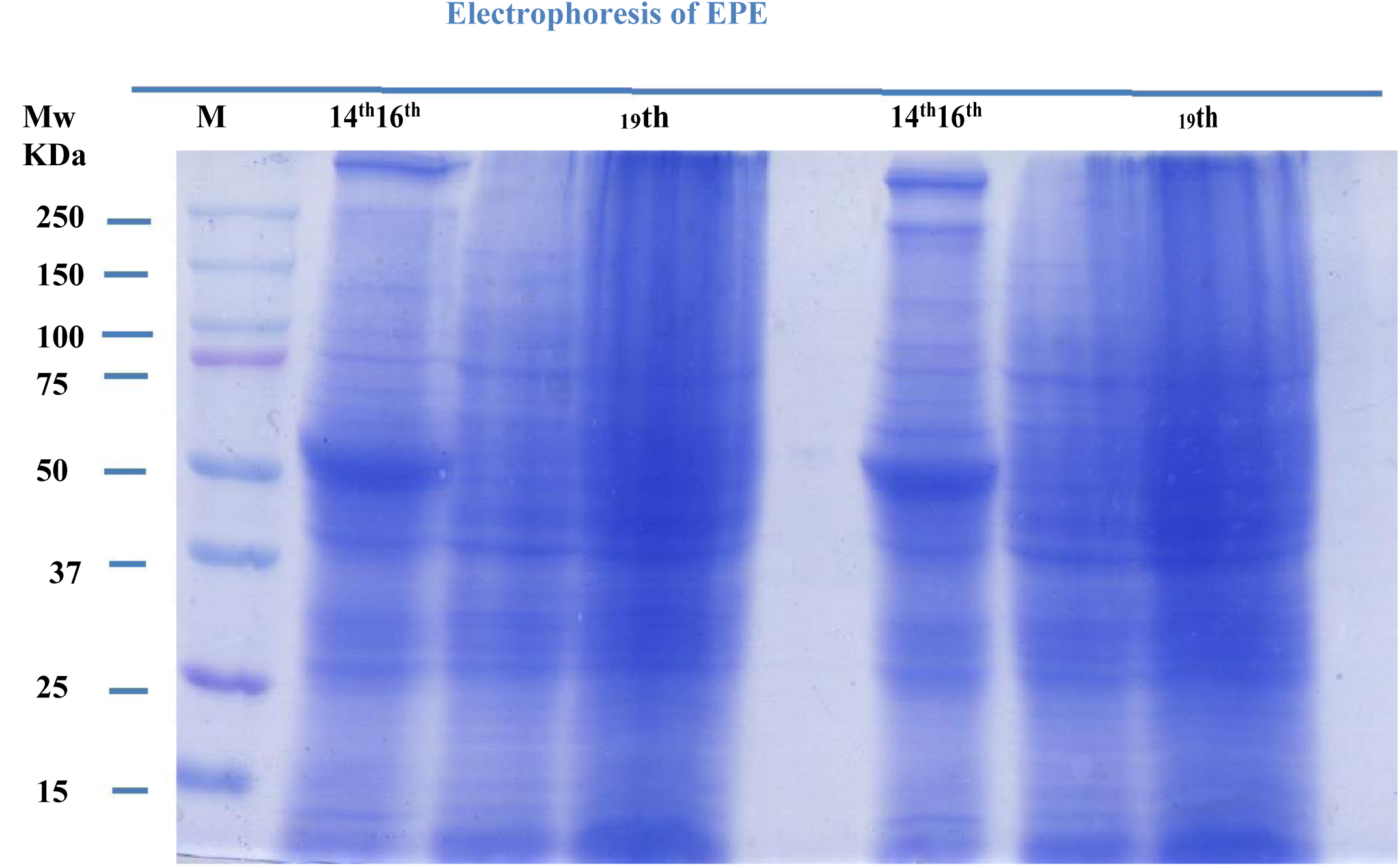
Proteins in the homogenized embryonic protein extract were separated on a 10% polyacrylamide gel. 1st well loaded with Biorad reference standard; 2nd, 3rd, and 4^th^ are loaded with 14th, 16th, and 19^th^ EPE; the same EPE samples were in duplicate.

### Injection of Embryonic Extract through Intravenous Route

Numerous in vitro studies have confirmed the biological functions of specific embryonic factors. Further studies are required to confirm *in vivo* regenerative function and safety [20]. In the present study we focused on the safety, so we have isolated embryonic protein and were injected through intravenously for *in vivo* safety evaluation [21]. 12 adult rats with an age of 13 weeks which were are divided into two groups, six rats of each were taken in control and treated groups for EPE injection. 120μL of EPE extract, which was prepared from 14^th^, 16^th^ and 19^th^ embryos days were injected to treated group of rats. Injected EPE has 0.724 mg/, 1.548 mg and 2.532 mg of protein confrontations respectively. After 48 hours of intravenous injection, blood sample was collected from both control and treated groups and tested for biochemical, hematological and immunological parameters [22, 23,24].

### Biochemical, Immunological and hematological parameters were tested

Serum Creatinine, blood urea, blood sugar, serum proteins, albumin and globulin are analyzed on fully automated biochemical analyzer Erba EM 200. Serum samples consumed for these tests are in the following sequence from creatinine to globulin, respectively: Creatinine 12 uL, urea 2 uL, sugar 2 uL, proteins 20 uL, albumin 2 uL. A total of 38 uL serum is utilized for biochemical Collected blood samples were analyzed to quantify biochemical parameters like serum creatinine analysis in each group. Graphical data presented in (Fig-4).

**Figure 4:**
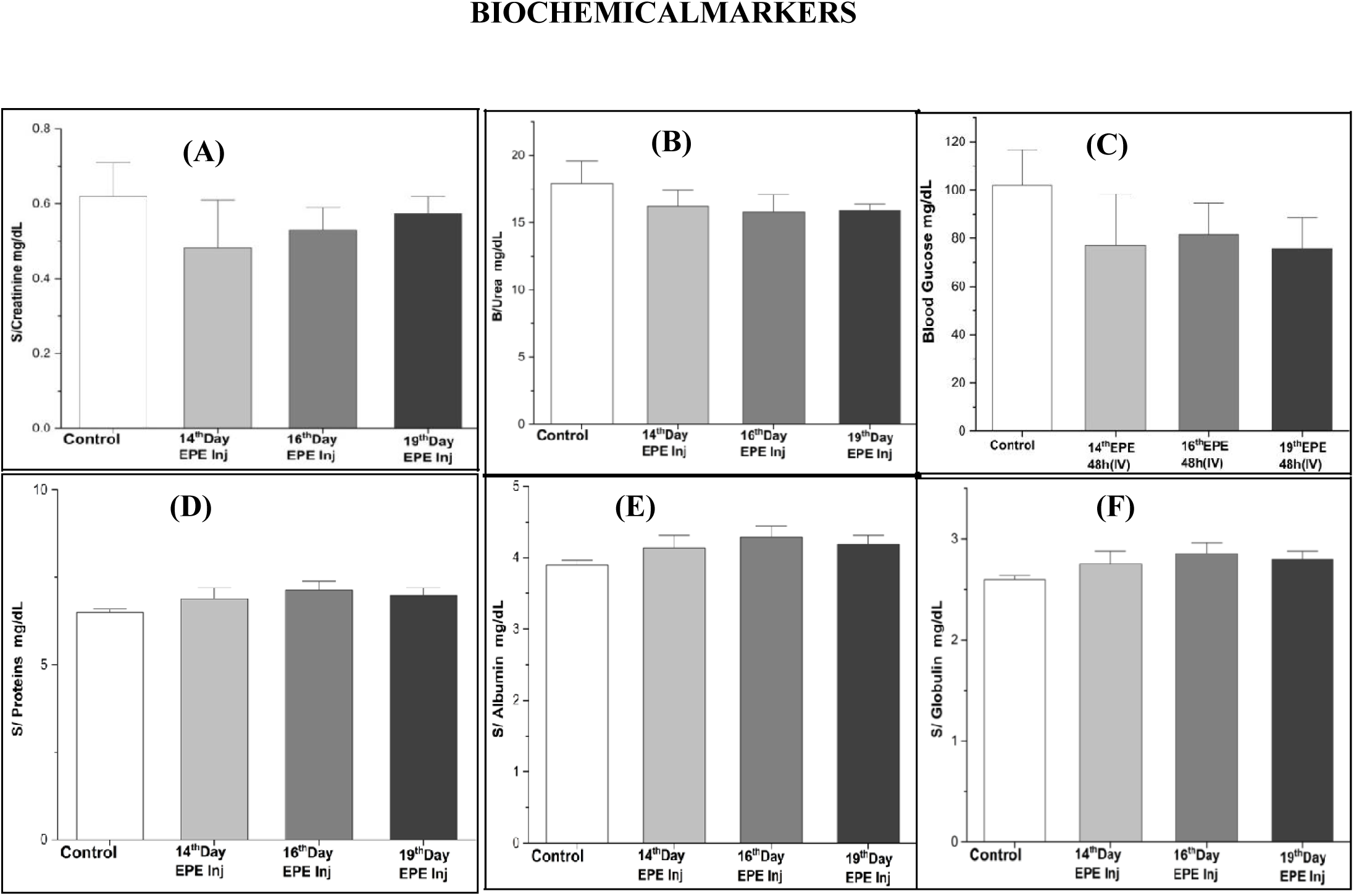
After 48 Hours Intravenous injection of embryonic extract, following biochemical parameters A) S/creatinine, B) B/urea, C)S/Proteins, D) S/Albumin, E) S/Globulin are tested and values are expressed as Standard error of the mean(mean ± SEM) table 1. Values are also plotted in bar graphs with standard error in each column includes control group, embryonic days 14^th^, 16^th^ and 19^th^ day embryonic extract injection. t-Test: Two-Sample Assuming Unequal Variances was also performed as mean ± SEM and P value for all parameters are significantly less then ( p < 0.005).

**Table 1:** The Mean values of (mean ± SE) of serum biochemical profile in control and treatment rats after 48 hours of intravenous injection of EPE.

| S.NO | PARAMETERS | ROUTE | CONTROL | 14<sup>th</sup> Day<br>48h | 16<sup>th</sup> Day<br>48h | 19<sup>th</sup> Day<br>48h |
| --- | --- | --- | --- | --- | --- | --- |
| 1 | CREATININE | IV | 0.62±0.0.25 | 0.48±0.038 | 0.53±0.02 | 0.57±0.021 |
| 2 | BLOOD UREA | IV | 17.93±0.052 | 16.2±0.037 | 15.7±0.29 | 15.9±0.24 |
| 3 | BLOOD<br>GLUCOSE | IV | 102.0± 4.8 | 77.0± 5.5 | 81.5± 3.7 | 75.8±3.3 |
| 4 | SERUM<br>PROTEINS | IV | 6.5±0.13 | 6.8±0.09 | 7.1±0.06 | 6.9±0.104 |
| 5 | ALBUMIN | IV | 3.9±0.08 | 4.1±0.05 | 4.3±0.04 | 4.2±0.06 |
| 6 | GLOBULIN | IV | 2.6±0.06 | 2.8±0.04 | 2.9±0.03 | 2.8±0.04 |

### Immunological Markers

CRP, IL6, levels were estimated by ELISA method 100 µL a microliter of blood was utilized. Elabscience kit was used for this analysis. ELISA studies of CRP and IL6 are in normal range, graphical data presented in (Fig. 5).

**Figure: 5.**
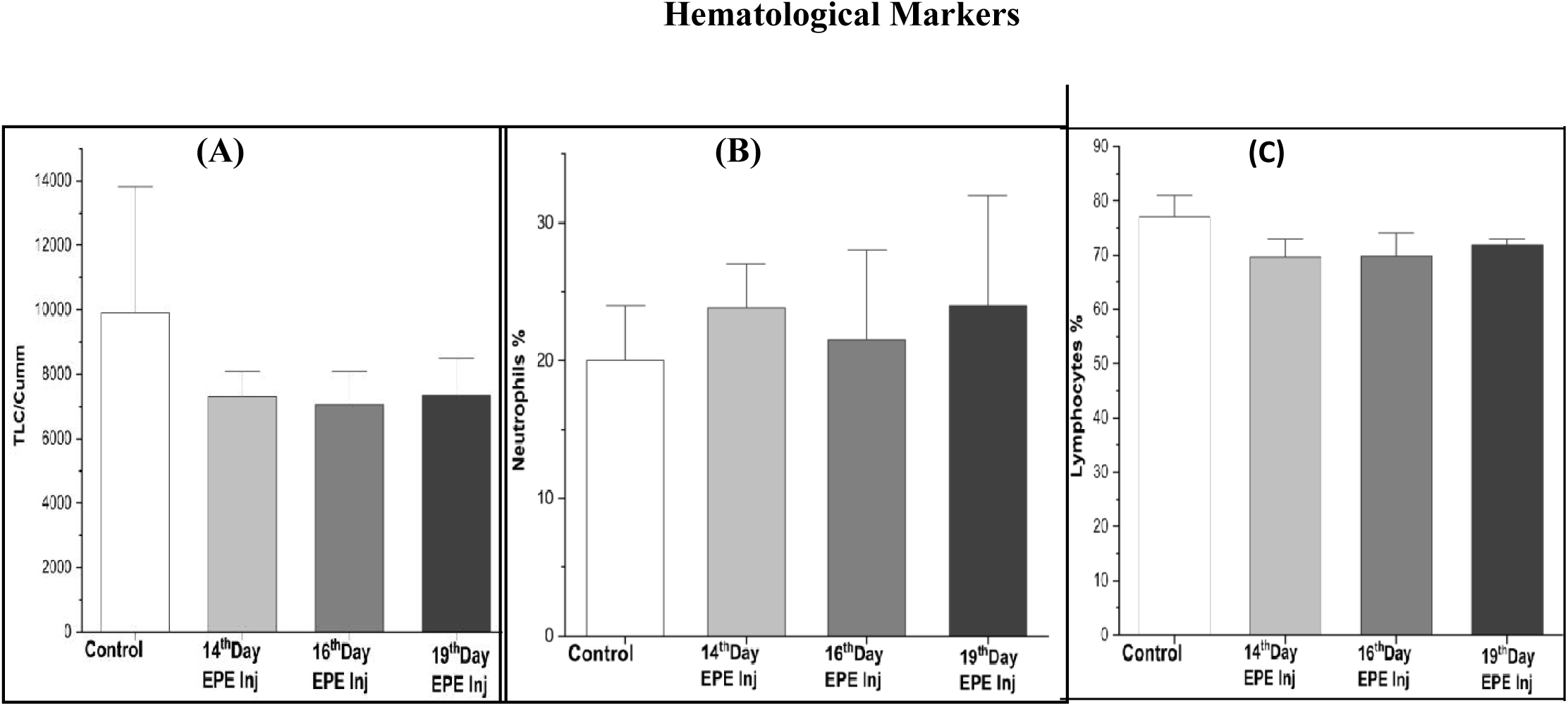
After 48 Hours Intravenous injection of embryonic extract, following haematology parameters (Total count, Neutrophils and Lymphocytes %) were performed and values are expressed as Standard error of the mean (mean ± SEM) table 2. Values are also plotted in bar graphs with standard error in each column includes control group, embryonic days 14^th^, 16^th^ and 19^th^ day embryonic extract injection. t-Test: Two-Sample Assuming Unequal Variances was also performed as mean ± SEM and P value for all parameters are less then ( p < 0.05).

**Table 2:** Mean and standard error values of hematological parameters after 48 hours of intravenous injection of 14^th^ 16^th^ and 19^th^ day of EPE to rats under control and treatment groups (n=6)

| S.No | Parameters | Control | 14<sup>th</sup> Day | 16<sup>th</sup> Day | 19<sup>th</sup> Day |
| --- | --- | --- | --- | --- | --- |
|  |  |  | 48h | 48h | 48h |
| 1 | **TLC** | 9,890±206.5 | 7,300±206.55 | 7,067±396.4 | 7,340±320.17 |
| 2 | **Neutrophils %** | 20±1.06 | 24±1.13 | 22±1.68 | 24±2.3 |
| 3 | **Lymphocytes %** | 77±1.18 | 69±1.97 | 70±1.68 | 72±2.02 |

**Table 3:** Mean and standard error values of CRP and IL6 at two different intervals 48 Hours of Intravenous injection of 14^th^ 16^th^ and 19^th^ day embryonic extract to rats under control and treatment groups (n=6)

| S.No | Parameters | Control | 14<sup>th</sup> Day | 16<sup>th</sup> Day | 19<sup>th</sup> Day |
| --- | --- | --- | --- | --- | --- |
|  |  |  | 48h | 48h | 48h |
| 1 | **CRP pg/mL** | 198.5±11.5 | 156.48±3.58 | 157.5±2.68 | 164.07±3.4 |
| 2 | **IL-6 pg/mL** | 19.5±0.8 | 17.1±0.38 | 16.78±0.41 | 16.67±0.28 |

### Haematological Parameters

TLC, neutrophils (%), and lymphocytes (%) were quantified. 250 µL of blood was collected in anti-coagulated tubes and was used to assess the haematological parameters [25], the total leucocyte count, percentage of neutrophils and lymphocytes are analysed on Cell counter Erba H 360, the graphical data presented in (Fig-6) These quantified parameters of both control and treated groups were compared through T-test, to find out significant difference

**Figure 6:**
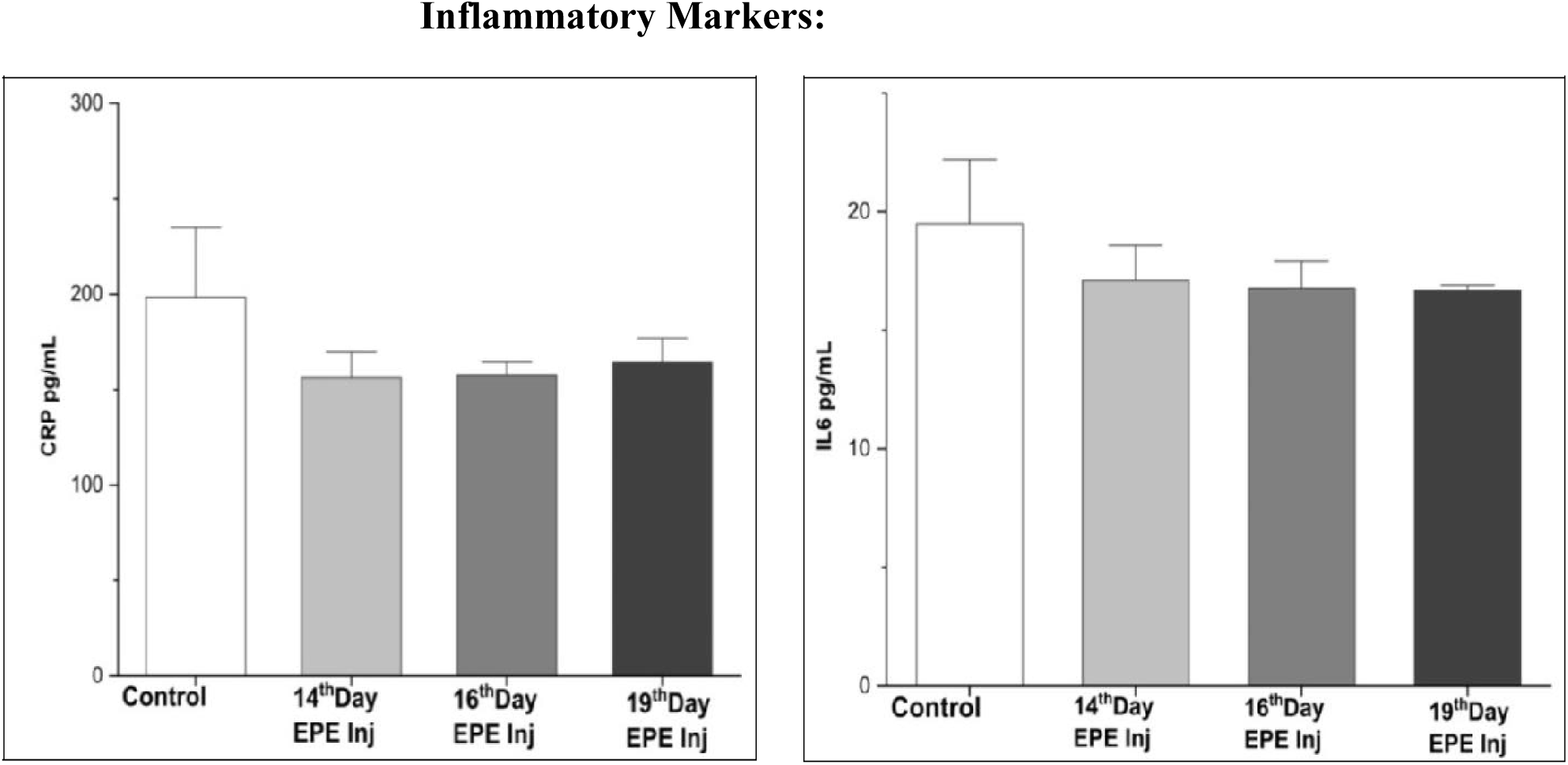
After 48 Hours Intravenous injection of embryonic extract, following immunological parameters (CRP and IL6) were performed and values are expressed as Standard error of the mean (mean ± SEM) table 2. Values are also plotted in bar graphs with standard error in each column includes control group, embryonic days 14^th^, 16^th^ and 19^th^ day embryonic extract injection. t-Test: Two-Sample Assuming Unequal Variances was also performed as mean ± SEM and P value for all parameters less then ( p < 0.05).

### H& E Staining

Histological examination procedures were followed as described by Adeyemi and Akanji[26]. Among six treated rats, one rats was euthanized and organs were collected rat as per (CCSEA), India guidelines. The brain, heart, kidney, large intestine, liver, small intestine, spleen and stomach samples were collected and fixed in 10% (v/ v) formaldehyde. Tissue samples were dehydrated through ascending grades of ethanol (70%, 90%, and 95%, v/v), then cleaned in xylene. The tissues were embedded in the paraffin wax (melting point 56^0^C) and stained with hematoxylin and eosin (H & E). The photomicrographs were captured at 100X using the software Presto Image Folio package (Fig. 7)

**Figure 7:**
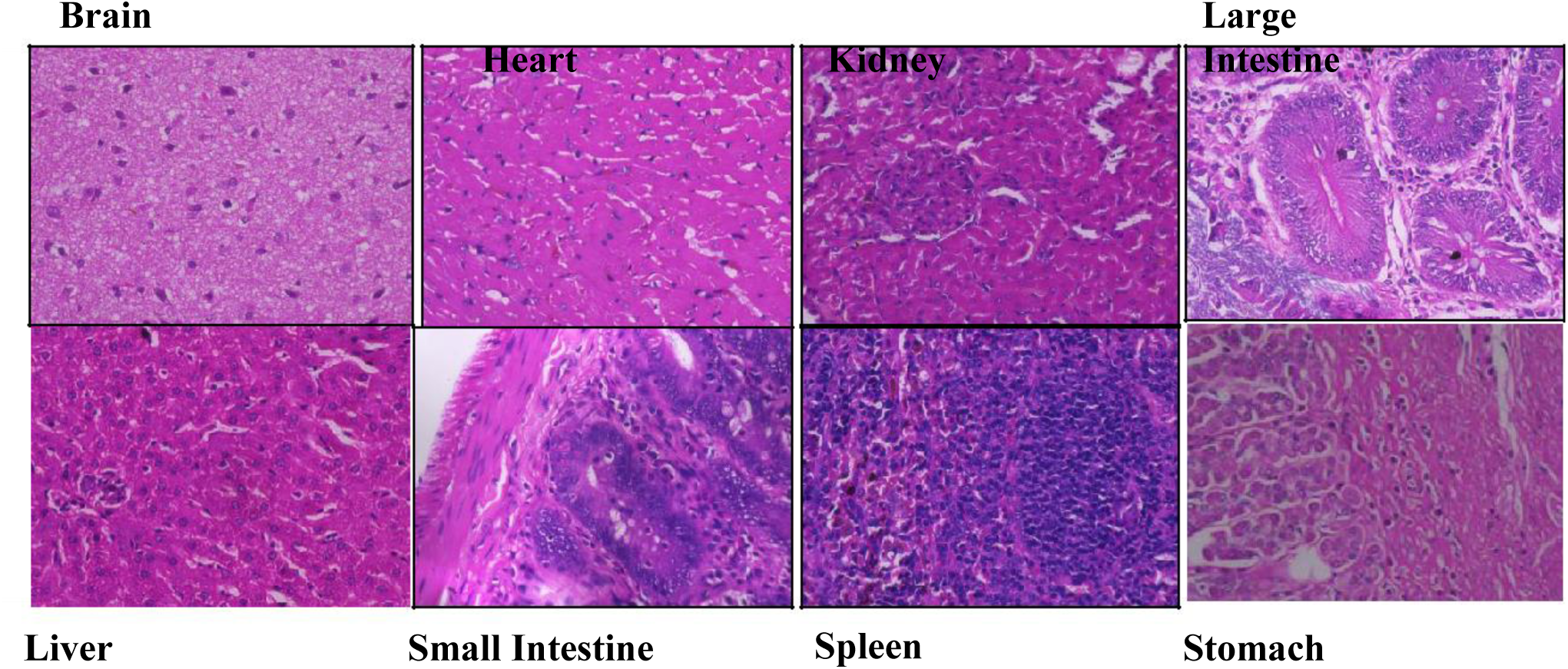
Histopathology study of tissues. Following EPE Intravenous injections, histopathology examination different tissues of treated rat was studied by staining with hematoxylin and eosin A) Brain, B) Heart, C) Kidney D) Liver, E) Small intestine and F) Large intestine, G) Spleen and H) stomach are with normal histoarchitecture without tarotamas

## RESULTS

### Biochemical Parameters

Biochemical parameters play a crucial role in drug testing. These parameters help in understanding the quantitative differences in various parameters, after intravenous injection of 14^th^ 16^th^ and 19^th^ day of EPE, the serum biochemical profile were estimated and compared the biochemical profile with control group values and all parameters are with in normal range. Graphical data of biochemical markers are presented in (Fig-4)

### Hematological Parameters

Hematological parameters can be affected by medications direct medication of EPE, including changes in leucogram and differential count especially neutrophils and lymphocytes, because EPE contains various growth factors that are not expressed in adult stage. Graphical data of Haematological markers are presented in (Fig-5)

### Immunological Markers

Vertebrate Immune system is more advanced and highly protected with immune system, if any foreign protein enters into the body, immune system responds instantaneously by synthesizing CRP from liver and IL6 by macrophages. In this study the estimated CRP and IL-6 are in normal range in both control and treatment groups. Graphical data of immune markers are presented in (Fig-6)

### Histological Examination of tissues After EPE injection

After EPE intravenous injection, results of the histopathological examination of rat brain, heart, kidney, liver, small intestine and large intestine of the treated rat and compared with normal organs of rat, showed no signs of teratomas or tumours in the tissue sections in above organs and cellular architecture in both treated and control groups are normal (Fig – 7). The data confirms that, EPE can be used for *in-vivo* for regenerative therapy in adult stag

### Statistical Analysis and Quantification

Bar overlap diagrams were generated using Origin software. The p-value was calculated by t-Test: Two-Sample Assuming Unequal Variances using Microsoft Excel indicated. All experiments (except the ones involving the animal models) were done at least with duplicates for each condition within the experiments. The data are represented as mean ± standard error, p-Value is less than 0.05.

## Discussion

In this work, we confirmed the safety of embryonic proteins by testing immunological, biochemical, and hematological markers after 48 hours of EPE injection. Biochemical parameters are crucial in vaccine studies as well as therapeutic protein injection studies. Intravenous injection of embryonic proteins is a new treatment methodology, so our biochemical parameters in this study can provide critical information about the acceptance, vaccine efficacy, and safety. In EPE intravenous injection studies, kidney and liver function parameters are considered; basic biochemical parameters like creatinine, urea, blood glucose, serum proteins, albumin, and globulins are quantified, and they are within normal limits.

Hematological parameters have been studied in various contexts, including vaccine studies and molecular therapy of proteins. EPE injection is also a new study, so we have studied fluctuations in TLC, the percentage of neutrophils and lymphocytes, after intravenous injection of EPE. Vertebrate’s immune system is sophisticated with innate immunity and adaptive immunity; both will respond instantaneously to foreign protein entry. In vivo application of EPE is a new approach, and we found no fluctuation in TLC, the percentage of neutrophils and lymphocytes, after EPE injection.

Inflammatory parameters are often synthesized when any non-self-protein enters the body. For this in vivo safety study of EPE, we have analyzed the CRP and IL6. Acute-phase proteins and inflammatory markers can help understand APRs and homeostatic changes. In this study, there is no absolute change in these parameters.

Tumorigeneciety and heterogeneity are major limitations in the use of stem cells in regenerative therapy. As tumorigeneciety and heterogeneity are cellular functions, we have deployed homogenization of embryonic stem cells to decellularize, so that embryonic cells are disrupted but molecules of embryonic cells are intact and are biologically active and preserved. After intravenous injections, HPE studies were also performed on different tissues to confirm that EPE injections are non-tumorigenic and non-heterogenic without the loss of cellular identity in tissues.

Studies **That Confirm the Biological Functions of Embryonic Proteins** Induced pluripotency is one of the classical examples of embryonic molecular function. The core transcriptional factors involved in the development of pluripotent cells are Oct3/4, Sox2, Klf4, and c-Myc. All these factors are called Yamanaka factors, named in honor of Noble Scientist Shenya Yamanaka. These factors were isolated from the 14^th^ mouse embryonic day [27].

In vitro neurogenesis, the process of generating new neurons in a laboratory setting, has been studied using various growth factors. Fibroblast growth factor 2 (FGF-2) has been shown to boost the production of neurons in human neural stem cell (hNSC) cultures [28].

Hepatocyte differentiation factors for in vitro culture have been investigated in several studies. A study identified a combination of four transcription factors (Hnf4a, Foxa1, Prox1, and Hlf) that promoted the differentiation of liver ductal organoids (LDOs) into mature hepatocyte-like cells with enhanced functionality [29, 30]. Another study found that hepatocyte growth factor (HGF) and activin A and Wnt3a were optimal for promoting the endodermal expansion of human umbilical cord mesenchymal stromal cells (HUCMSCs) into hepatocyte-like cells (HLCs). [31]

Molecular nephrogenesis, the process of embryonic kidney development with the involvement of Cbx1, Cbx3, Cbx5, and Trim28, was confirmed by proteomic analysis of rat embryonic kidneys at different developmental stages [32]. Researchers in this study aim to identify proteins involved in the epigenetic regulation of gene silencing and their potential role in nephrogenesis. The embryonic kidney develops through three stages: pronephros, mesonephros, and metanephros. The metanephros forms the final kidney [33].

Integration of this present safety study and previous in vitro embryonic molecular function studies helps in developing organ-specific molecular regenerative therapy as well as a new arena of molecular therapy for reversing biological aging.

### Strengths and limitations

Direct injection of embryonic factors is possible with our study, as biochemical, immunological, and hematological parameters are within normal limits. We need to extend our study for proteomic analysis of embryonic protein extracts and confirmation of embryonic proteins in organ function restoration. For future applications in regenerative therapy, there is no need for scarification of embryos, as bulk industrial production can be feasible using biotechnological tools, and it should be carried out on a large group of experimental animals. Simultaneously, we are continuing this study, in which we injected embryonic protein extract through ultrasound-guided intra-renal injection. With this, we have confirmed the safety of intra-organ injections. After this injection, renal parameters came back to normal in the drug-induced chronic kidney disease (CKD) model. A proteomic study and a histopathology study of recovered kidneys are under analysis.

## Conclusion

Our study has ratified the safety of embryonic proteins by direct intravenous injection. Our study laid the foundation for future embryonic molecular regenerative therapy and also provided alternatives for stem cell-based therapy complications like heterogeneity, immunogenicity, and tumorigenicity, as embryonic proteins are non-immunogenic, non-tumorigenic, and non-heterogenic without altering cellular identify. With this study molecular based regenerative therapy replaces Stem cell-based Regenerative therapy.

## Ethical committee approval

Entire study was approved by Institutional Animal Ethical Committee (IAEC) of Jeeva Life Science with the approval number: CPCSA/IAEC/JLS/13/08/20/04.

## Acknowledgment

I would like to thanks different research centers supported this work, AKP Nephrolab Department of Biochemistry, University of Hyderabad, Veterinary Pharmacology & Toxicoloyg and Veterinary Biochemistry, PV Narsimha Rao Telangana Veterinary University, Hyderabad, Telangana, Department of Physiology, Apollo Institute of Medical science and research, Hyderabad, Kidney and Laparoscopic centre, Hyderabad Blood Centre and Satya charitable Polyclinic, Hyderabad.

